# Visual dependence in postural control is increased in older adults

**DOI:** 10.1101/2025.08.06.668916

**Authors:** Saskia Neumann, Cyrille Mvomo, Deepak K. Ravi, Friederike A. Schulte, Lorenz Assländer, Chris Easthope Awai

## Abstract

Successful postural control depends on the integration of visual, vestibular, and proprioceptive inputs. With age, postural control degrades, leading to impaired balance and greater fall risk. Understanding how this integration changes over the lifespan is invaluable for designing more effective interventions that enable healthy postural control in older age. Earlier studies measured visual dependence using perceptual tasks or spontaneous sway comparisons across visual conditions. This study evaluates how visual dependence differs between younger and older adults within the postural control mechanism using a Central Sensorimotor Integration (CSMI) test. Eighty healthy adults (n=40, 60-87 years, n=40, 21-52 years) were exposed to small pseudorandom visual scene movements implemented in virtual reality while standing on a compliant surface. Sway responses were measured using virtual reality trackers and interpreted using an established frequency domain balance control model. Model parameters included visual weight, proportional and derivative feedback gains, time delay, and torque feedback gain. Test–retest reliability was assessed in a subgroup (n = 40) and showed excellent intra-class correlation coefficients for visual weight, proportional and derivative feedback gains (ICC = 0.89– 0.96), and lower ICCs for time delay (ICC=0.59) and torque parameters (ICC=0.39). The main difference between age groups was visual dependence, with older adults relying 40% on vision, compared to 33% for the younger group (p = 0.042). No significant group differences were found in other model parameters. Our results provide direct evidence of an increase in visual contribution to posture control with age.

## Introduction

The central nervous system’s ability to perceive, process, and integrate multisensory information changes over the lifespan and is affected by aging (Murray et al., 2016). Among the sensory systems, vision often dominates over vestibular and proprioceptive input, influencing postural control even when other cues may be more appropriate (Horiuchi et al., 2021). This phenomenon, known as visual dependence, refers to an excessive or disproportionate reliance on visual information for spatial orientation and postural control, even in situations where vestibular or proprioceptive cues would be more reliable (Maire et al., 2017). Several studies have provided indications that visual dependence increases over the lifespan and is linked to a greater risk of falling (Barr et al., 2016; Lord & Webster, 1990). This association is particularly important because falls are a leading cause of injury, loss of independence, and mortality among older adults (Rubenstein, 2006). An increased reliance on visual input may reflect age-related changes in the multisensory balance system. Understanding how visual weighting shifts with age is therefore crucial, not only to identify early signs of impaired balance control, but also to inform age-specific interventions aimed at preserving postural stability and functional independence.

Vision plays a key role in postural control by providing external spatial reference cues that help the central nervous system estimate body position and orientation in space (D. N. Lee & Lishman, 1975). With advancing age, visual function declines, which is characterized by reduced acuity, decreased contrast sensitivity, narrowing of visual fields, and slower dark adaptation (Salvi et al., 2006). These age-related changes can impair multiple tasks of daily life and contribute to reduced mobility and physical activity, which in turn impact broader aspects of health and independence (Swenor et al., 2020). Despite the well-documented decline in visual function, older adults are often reported to rely more heavily on visual input for maintaining balance compared to younger individuals (Bednarczuk et al., 2021; Osoba et al., 2019). This increased visual dependence may impair postural control, particularly in complex real-world environments with poor, misleading, or conflicting visual information. Visual dependence has been identified as both an early marker of balance impairment and an independent risk factor for falls (Barr et al., 2016; Bednarczuk et al., 2021; Danna-dos-Santos et al., 2021; Lord & Webster, 1990). One likely explanation for this increased reliance on vision is that it serves as a compensatory strategy in response to age-related declines in other sensory systems, even in absence of pathology. These cumulative sensory changes can further compromise postural stability and functional independence (Anson & Jeka, 2016; Henry & Baudry, 2019a).

Human standing balance is widely described as a feedback-driven, closed-loop system in which multiple subsystems (e.g. sensory systems, sensory integration processes, neuromuscular control and the mechanical properties of the body and musculotendinous structures) interact with one another and contribute to the behavior of the entire system (Peterka et al., 2018). This complex interplay complicates efforts to attribute balance impairments to specific sensory deficits or other changes in the system, such as time delays or feedback gains (Peterka et al., 2018). Assessing the contribution of the visual system in postural control has potential for identifying individuals at increased risk of falls and informing and guiding the development of targeted intervention strategies. Various methods have been developed to assess the visual dependence, but past work has primarily relied on indirect measures of visual dependence such as perceptual tests like the ‘Rod and Frame Test’ (Witkin & Asch, 1948) and the ‘Rod and Disk Test’ (Dichgans et al., 1972). These tests measure the influence of misleading visual cues on an individual’s sense of verticality. Older adults typically show greater susceptibility to such visual distortions compared to younger individuals, indicating an increased reliance on visual input for spatial orientation (S.-C. Lee, 2017). However, a key limitation of these tests is that they assess perceptual orientation and do not provide direct insights into the use of vision in postural control. Other common tests of visual dependence compare spontaneous sway under different visual conditions, typically between eyes-open and eyes-closed. However, spontaneous sway assessments have inherent limitations, as they do not differentiate between biomechanical, sensory and motor contributions to balance and are influenced by both internal noise and the feedback dynamics of the control system (Pasma et al., 2014). Consequently, spontaneous sway lacks the specificity needed to isolate the effect of changes in visual dependence from other subsystems (Peterka et al., 2018).

To assess visual dependence in postural control, externally applied balance perturbations and appropriate analytical models can be used to accurately disentangle the complex relationship between biomechanical, sensory, and motor contributions in a closed-loop system (Van Der Kooij et al., 2005). The most prominent test in the field eliciting these methods is the Central Sensory-Motor Integration (CSMI) test (Peterka, 2002; Peterka et al., 2018). The CSMI approach utilizes sensory perturbations to probe sensory contributions and characterize the balance control mechanism in the form of frequency response functions (FRFs) (Peterka, 2018). This approach enables the dissociation of internal noise from the underlying control dynamics, allowing for model-based interpretations to separate sensory, motor, and biomechanical contributions. These FRFs characterize postural behavior and are interpreted using a model-based parameterization. The identified parameters are physiologically interpretable, such as sensory weights, neural feedback gains, and time delays. This makes the CSMI test a powerful tool for quantifying sensory contributions in balance control.

To separate the visual contribution from the vestibular and proprioceptive system, we applied a virtual reality (VR-)based version of the CSMI test to compare visual dependence in postural control during standing between younger and older adults. We hypothesized that older adults would demonstrate higher visual weighting than younger adults. To further evaluate the robustness of this method, we conducted a test–retest reliability analysis in a subgroup of participants.

## Materials and Methods

### Participants

This study included 40 younger and middle-aged (aged >18 and <60 years) and 40 older (> 60 years) able-bodied adults. Eligibility criteria were community-dwelling individuals without any diagnosed acute or chronic musculoskeletal conditions (e.g., osteoporosis, rheumatoid arthritis, osteoarthritis) or neurological disorders (e.g., Parkinson’s disease, Multiple Sclerosis, Stroke). Participants were excluded if they reported difficulties in walking, were taking anti-depressant medications such as benzodiazepines, or had received artificial joint replacements within the last 1.5 years. Participants were recruited via flyers distributed in local community centers, at events, and sports clubs across the Canton of Lucerne, Switzerland. The study received ethical approval from the Ethics Committee of the Eidgenössische Technische Hochschule (ETH) Zurich (protocol number 2021-N-90), and all participants provided written informed consent prior to participation.

### Experimental setup

A schematic representation of the experimental setup is shown in Figure 1. Participants wore a VR head-mounted display (HMD; Vive Pro 2, HTC, Taoyuan, Taiwan) featuring a horizontal field of view of 116 degrees, a vertical field of view of 96 degrees and a resolution of 4896×2448 pixels. Motion tracking was performed using two diagonally placed base stations (Base Station 3.0, HTC, Taoyuan, Taiwan) and two Vive 3.0 trackers (HTC, Taoyuan, Taiwan) attached with Velcro straps at the lower back (L5) and between the shoulder blades. To minimize auditory cues and reduce distraction from the balance task, participants wore noise-canceling over-ear headphones (Sony WH-1000XM4) and listened to an audiobook. Participants stood on a foam pad (AIREX Balance Pad, ch_93019) to reduce the proprioceptive contribution and to increase the sensitivity to the visual perturbation (Schut et al., 2017). In the virtual environment, participants viewed a half-cylindrical screen with a radius of one meter and a high-contrast pattern (Assländer et al., 2023). During trials, the screen tilted in the anterior-posterior direction around the ankle joint axis. The tilt sequence consisted of 11 successive repetitions of a 20-s long pseudorandom sequence with a peak-to-peak amplitude of 1.2 ° corresponding to peak velocities of ±0.5 °/s (Peterka, 2018).

**Figure 1.**
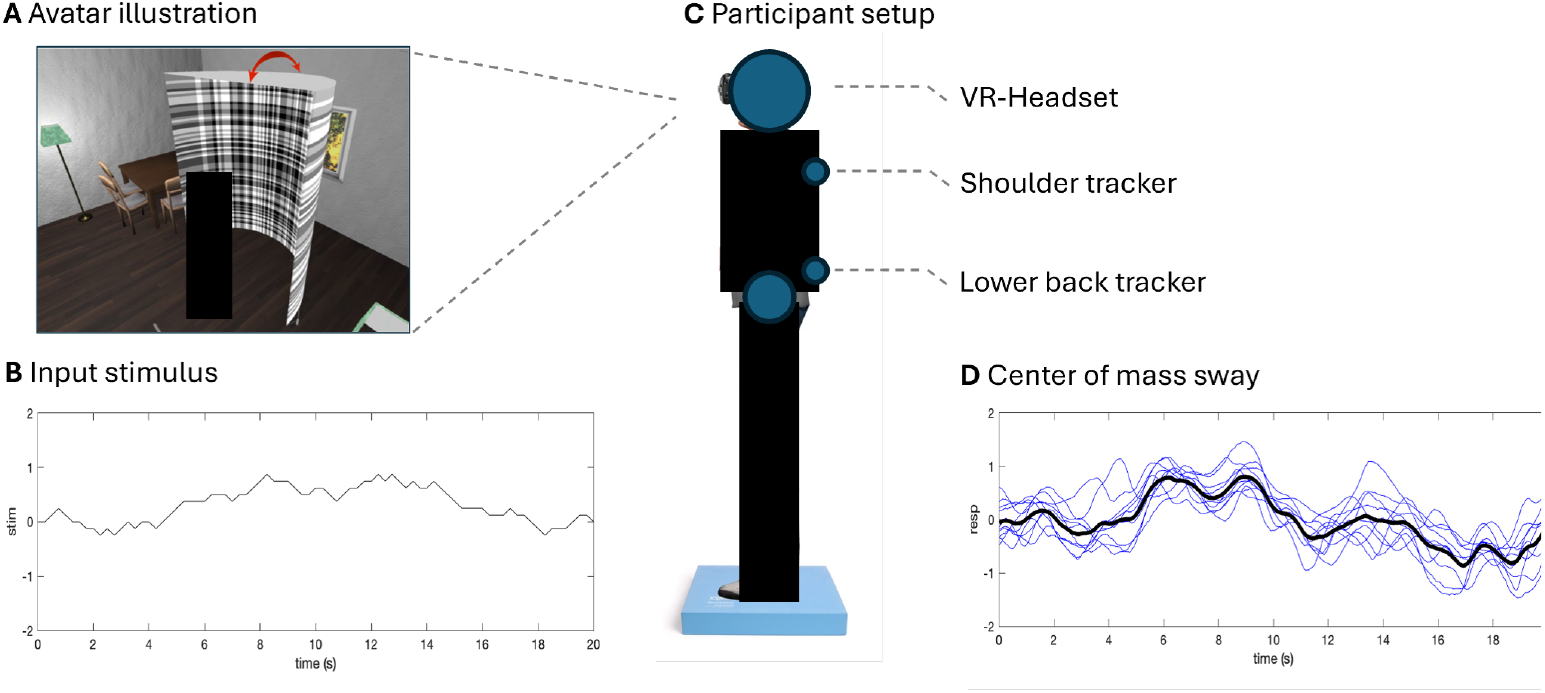
Experimental setup. **A Avatar illustration** of the participant’s position within the virtual environment; **B Input stimulus** which was repeated 11 successive times; **C Participant setup** with VR-headset, shoulder and lower back tracker while standing on a foam pad; **D Center of mass sway** of the subject in individual stimulus repetitions (blue lines) and the average across repetitions, representing an approximation of the response to the stimulus.

### Procedures

After providing informed consent, participants’ weight and height were recorded. CSMI measurements were conducted as pre- and post-tests of a randomized controlled trial evaluating the effects of a single-session perturbation-based balance training, which included both an intervention and a control group for each age group. As the pre-test (T1) was used for the main analysis, the perturbation training had no influence on the presented results. In addition, we used the post-test data (T2) from the control group (n = 20 per age group) to assess the test-retest reliability of the CSMI measurement. The post-test was administered a second time following a 24-minute treadmill walking session, which served as a control condition. All participants were naïve to the test procedures to ensure unbiased responses. To allow familiarization with the virtual environment, participants wore the VR-headset and stood on the foam pad for 1–2 minutes before testing began. During the experiment, participants wore shoes, were instructed to remain silent and encouraged to stand independently in a comfortable position on the foam pad, with feet hip-width apart and arms relaxed at their sides.

### Data analysis

The CSMI test is described in detail elsewhere (Assländer et al., 2023; Peterka, 2018). In brief, we calculated center of mass sway from hip and shoulder tracker movement. The first stimulus cycle was omitted to avoid transient responses and the remaining ten were averaged and used as approximation of the response to the stimulus. Sway response power was calculated as a descriptive measure for the amount of sway evoked by the stimulus, using the mean of the squared sway around the average body position. Random sway power was obtained by calculating the mean across time of the variance across stimulus cycles (see remnant sway calculation in Van Der Kooij & Peterka, 2011).

Average sway response spectra were further used to calculate the FRF, by dividing the center-of-mass sway spectrum by the stimulus spectrum at all frequencies with non-zero stimulus energy. We fit a transfer function formulation of the balance control model to the FRF of each individual participant, adjusting five parameters: Visual weighting (W_v_), proportional feedback gain (K_p_), derivative feedback gain (K_d_), time delay (T_d_) and the contribution from a low-pass filtered positive torque-feedback loop (K_t_). Body inertia J and gravitational forces mgh were calculated from participant body weight and height using anthropometric tables (David A. Winter, 2009). The transfer function is given by:

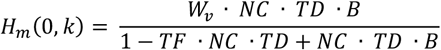

with the linearized (small angle approximation sin(γ) ≈ γ), the body dynamics (B = (J s^2^ - mgh)^−1^), the neural controller (NC = Kp + s Kd), the time delay (TD = exp^−s Td^), the low-pass torque feedback (TF = Kt s^−1^) and the Laplace variable s(*k*) = jω(*k*) for frequencies k. All analyses of sway data were performed using Matlab and the global optimization toolbox (The Mathworks, Natick, USA). Further details can be found in Assländer et al. (2023).

W_v_, represents the proportion of the overall torque that is attributed to visual inputs in the process of sensory integration for maintaining balance. Essentially, W_v_ quantifies the contribution of visual information to the control of balance, expressed as a percentage. The remaining contribution, which is 100% − W_v_, comes from the proprioceptive and vestibular inputs. These non-visual contributions are combined and cannot be separated without additional perturbation techniques. The parameters K_p_ and K_d_ represent the proportional and derivative feedback gains, respectively. The feedback time delay parameter T_d_ encompasses all time delay components within the control mechanism, such as neural conduction times and muscle activation delays. Lastly, K_t_ represents the contribution from a low-pass filtered positive torque-feedback loop, proposed to explain low-frequency sway characteristics with a period > 20s. Model parameters were estimated individually using an optimization procedure that minimized the error between the experimental and simulated FRFs. Specifically, a Maximum Likelihood−based optimization algorithm was used to adjust model parameters, aiming to reduce the discrepancy between the measured and predicted FRFs. The model formulation and parameter estimation followed the methodology described by Peterka et al. (2018) and Assländer et al. (2023). The output parameter description is summarized in table 1.

**Table 1.** CSMI output parameter description.

| Parameter | Description |
| --- | --- |
| $W_v$ | Visual weight, the percentage of the visual contribution to the overall torque. |
| $K_p$ | Proportional feedback gain, strength of the muscle contraction in response to deviations from the desired upright position (feedback error). |
| $K_d$ | Derivative feedback gain, strength of muscle contraction relative to the velocity of the feedback error. |
| $T_d$ | Lumped feedback time delay. |
| $K_t$ | Contribution from a low-pass filtered positive torque-feedback loop. |

### Statistical analysis

All statistical analyses were conducted using RStudio (Version 2024.09.0+375). Prior to analysis, the normality of continuous variables was assessed using the Shapiro-Wilk test. For parameters that were normally distributed, independent samples t-tests were used. For non-normally distributed parameters, the Mann–Whitney U test was applied. Effect sizes were calculated using Cohen’s d (d) for normally distributed data and rank-biserial correlation (r) for non-normally distributed data. According to Cohen (2013), values of d ≤ 0.2 indicate a small effect, 0.2 ≤ d ≤ 0.5 indicate a medium effect, and d ≥ 0.8 indicate a large effect. For a rank-biserial correlation, values of r ≤ 0.3 indicate a small effect, 0.3 ≤ r ≤ 0.5 indicate a medium effect, and r ≥ 0.5 indicate a large effect (Kerby, 2014). Statistical significance was determined at a threshold of p < 0.05. To control for multiple comparisons across the five model output parameters, the Simes correction was applied to adjust p-values. Outliers were defined as values exceeding ± 2 standard deviations from the group mean and were excluded from analysis. To assess the test-retest reliability of the CSMI parameters, the intraclass correlation coefficient (ICC 3,1) and corresponding 95% confidence intervals were calculated to evaluate agreement between the two measurement timepoints (T1 = pre-test 1, T2 = post-test). Based on the guidelines by Koo and Li (2016), ICC values <0.50 indicate poor reliability, 0.50 ≤ ICC < 0.75 indicate moderate reliability, 0.75 ≤ ICC < 0.90 indicate good reliability, and values >0.90 indicate excellent reliability.

## Results

All participants completed the experiment with n = 40 younger and n = 40 older participants. No adverse events occurred during testing. One participant from the younger group was excluded from the analysis as their data exceeded two standard deviations from the group mean and was therefore considered an outlier. A comparison of the general characteristics between the two age groups revealed no significant differences, see table 2. Descriptive response sway power (z = −2.18; *p* = 0.029, *r* = 0.29) and random sway power (z = 3.73, *p* < 0.001, *r* = 0.49) showed significant differences between the groups with higher values for the older group. The parameter W_v_, representing visual weight differed significantly between the age groups, *t*(77) = 2.71, *p* = 0.042) indicating a medium effect size (*d* = −0.61). The mean visual weight was 32.7 ± 10.9% in the younger group and 40.0 ± 12.9% in the older group, with a mean difference of 7.3% (95% CI [1.9%, 12.6%]). The group difference analysis was repeated including the outlier participant. After correcting for multiple comparisons, the group difference was no longer statistically significant (t(78) = 2.29, *p* = 0.088, *d* = −0.51). All other model parameters did not show significant differences between the groups with K_p_ (z = −1.75, *p* = 0.200, *r* = 0.23), K_d_ (z = −0.86, *p* = 0.646, *r* = 0.11), T_d_ = t(77) = 0.31, *p* = 0.950, *d* = −0.07) and K_t_ (z = −0.25, p = 0.810, r = 0.03). Figure 2 presents the group differences for all model parameters and the descriptive parameters and their distributions and individual data points. In addition to the visual representation, the figure also includes the corresponding statistical comparison.

**Table 2.** Participant characteristics of participants groups.

|  | Younger adults | Older adults | *p* |
| --- | --- | --- | --- |
| Number of participants | 39 | 40 |  |
| Gender (M/F) | (21/18) | (20/20) |  |
| Age (years, mean ± STD) | 29.69 ± 8.78 | 71.18 ± 6.36 | $< 0.001$ |
| Height (m, mean ± STD) | 1.71 ± 0.09 | 1.68 ± 0.07 | 0.120 |
| Weight (kg, mean ± STD) | 69.55 ± 11.72 | 75.67 ± 15.37 | 0.051 |

**Figure 2.**
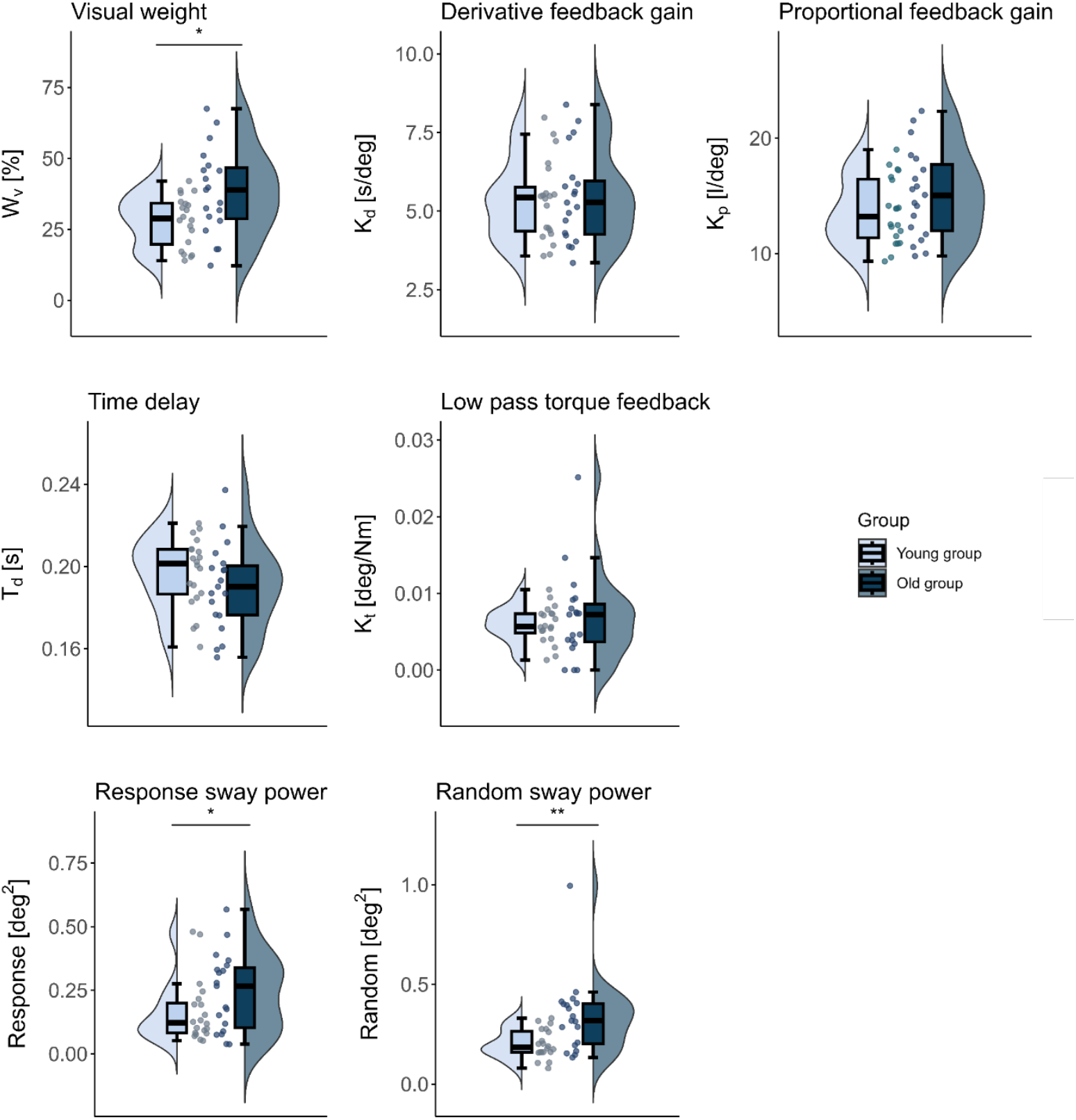
CSMI parameters and descriptive sway power values for the mean across all cycles (response sway power) and the variance across cycles (random sway power). */** indicate statistically significant group differences between younger and older adults (*p<0.05, **p<0.01).

The virtual CSMI test was performed twice for a subgroup of 40 participants, 20 per age group, to assess the test-retest-reliability of the parameter estimates. Figure 3 presents the CSMI parameters, simulation error, response sway power, and random sway power, along with corresponding ICC(3,1) values and their 95% CI. Parameter estimates showed large within and between subject variability. ICC values ranged from 0.301 to 0.966, indicating varying levels of reliability across parameters.

**Figure 3.**
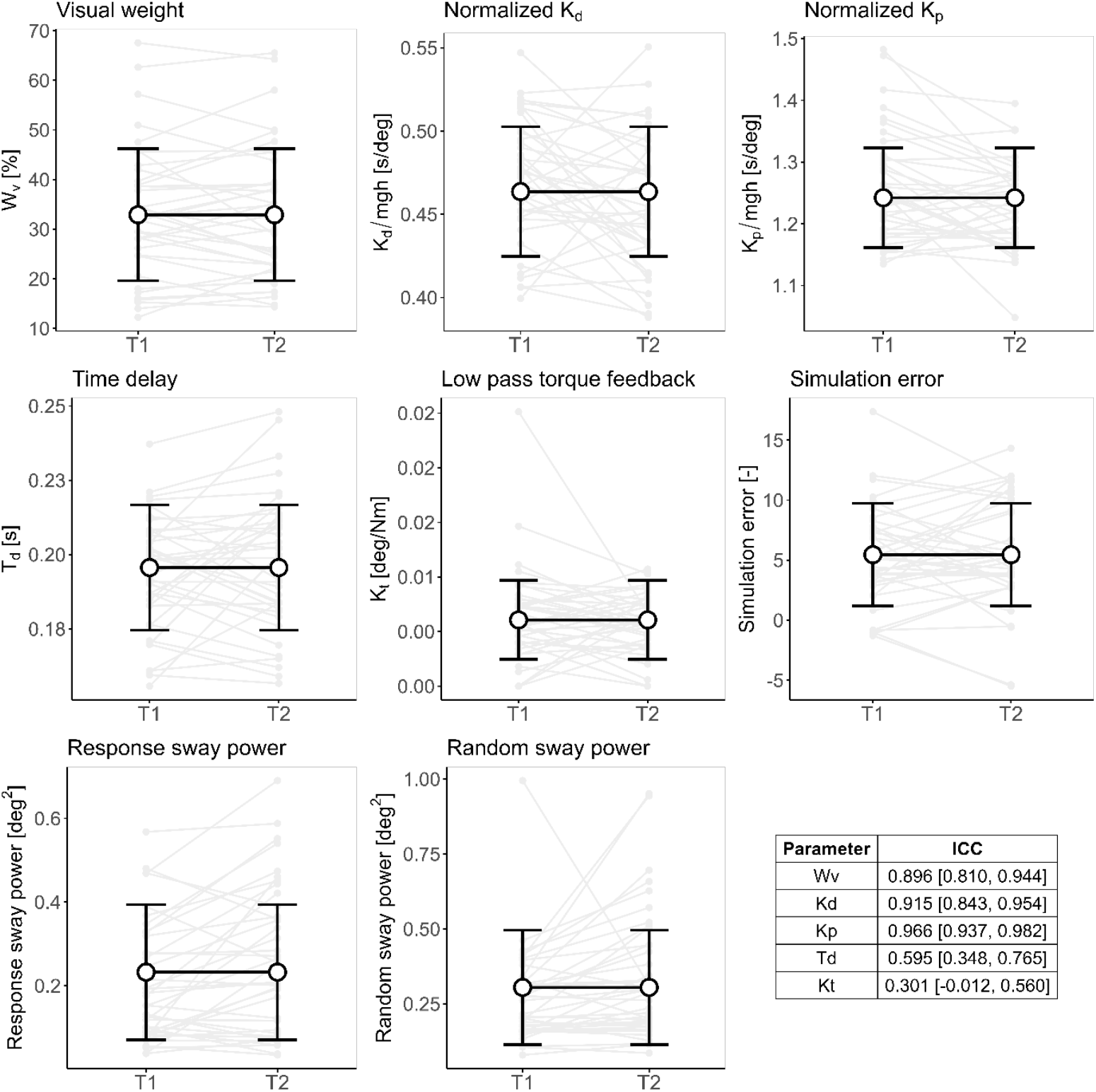
CSMI parameters for both timepoints (T1 and T2) for the subgroup of participants. Individual subject data are shown in light grey, along with the across-subject mean, and standard deviation. K_p_ and K_d_ are normalized by mgh. Intraclass correlation coefficients (ICC(3,1)) with 95% confidence intervals are presented in the accompanying table.

## Discussion

This study showed that visual dependence in multisensory integration underlying balance control is increased in older adults compared to younger adults. By comparing outcomes between these two groups, this study reports age-related differences in visual dependence for postural control. As hypothesized, older adults exhibited greater visual dependence for postural control compared to younger adults. Additionally, a subgroup analysis assessed the test-retest reliability of CSMI parameter estimates.

The CSMI test while standing on foam revealed that older adults generate 40% of overall torque based on vision compared to 33% in younger adults. As an increase in the weighting of vision necessarily corresponds to a decrease in the relative contribution of the other sensory modalities, the proprioceptive and/or vestibular contributions are reduced in older adults. The increased visual dependence observed in older adults aligns with previous studies using alternative methods, such as spontaneous sway analysis under eyes-open and eyes-closed conditions (Bednarczuk et al., 2021) and from perceptual studies (S.-C. Lee, 2017; Lord & Webster, 1990). These findings are also consistent with the model proposed by Henry and Baudry (2019b), which suggests that declining proprioceptive reliability with age leads to compensatory upweighting of visual input during balance control. This sensory adaptation is often accompanied by increased co-contraction of lower limb antagonist muscles and greater cognitive involvement. While these strategies may partially offset sensory deficits, they are also associated with reduced efficiency and adaptability of the postural control system in older adults. Apart from visual weighting, none of the other CSMI model parameters differed between the age groups. This suggests that sensory integration is a major factor in age related deteriorations, while other aspects are less affected. Thus, our results are overall in agreement with previous findings, but expand current knowledge through the specificity of the CSMI test.

The CSMI parameters provide a detailed interpretation of the changes in sway response power. However, the model-based interpretation does not address changes in random sway. This sway component reflects spontaneous postural fluctuations that are not evoked by the visual scene movement. The elevated random sway levels found in our study are consistent with prior reports of age-related increases in mean sway velocity and frequency (Maki et al., 1990; Maurer & Peterka, 2005; Qu et al., 2009), likely reflecting age-related declines in sensory and motor control (Sullivan et al., 2009). For example, older adults tend to exhibit higher center-of-pressure velocity and larger sway areas during simple balance tasks, such as the Romberg test, when compared to younger individuals (Bergamin et al., 2014). Notably, increased postural sway during quiet standing has also been identified as a strong, independent predictor of future falls, even after controlling for multiple confounders (Johansson et al., 2017). The elevated random sway seen in older adults may reflect compensatory adjustments in response to sensory decline, whereby the balance system relies on higher-amplitude corrective movements to maintain stability. However, this compensation may be only partially effective, as excessive sway variability is often associated with reduced postural control efficiency and increased fall risk (Kanekar & Aruin, 2014).

The sub-group analysis demonstrated high test-retest reliability for the parameters W_v_, K_d_ and K_p_, with ICCs of 0.896, 0.915, and 0.966, respectively. In contrast, the parameters T_d_ and K_t_ demonstrated poor and moderate reliability, respectively, indicating that temporal dynamics and torque feedback measures were less consistent across sessions compared to the other model parameters. In a previous study, test-retest reliability was evaluated using a comparable experimental setup, involving six repetitions of a 6 × 20-second pseudo-random sequence of visual scene tilts across three repetition per day in 14 young, healthy participants using a VR application (Assländer et al., 2023). In that study, similar ICC values were reported for W_v_, K_d_, and K_p_ (ranging from 0.70 to 0.92), supporting the robustness of these parameter estimates across different methods and participant samples. Compared to Assländer et al. (2023), the present study showed lower reliability for T_d_ and K_t_, which may reflect greater interindividual variability or reduced stability in the temporal characteristics of postural responses. The wider confidence intervals for these parameters further highlight this variability, suggesting that temporal aspects of balance control are more susceptible to external influences or may be less reliably captured by the current experimental setup. One likely contributor to the reduced reliability of T_d_ in this study is the use of a compliant foam pad, which was not employed in the Assländer et al. (2023) protocol. Compliant surfaces, such as foam pads, are known to degrade proprioceptive feedback, thereby increasing reliance on vestibular and visual cues (Shumway-Cook & Horak, 1986). While such surfaces are valuable for identifying subtle balance deficits by challenging proprioceptive input, the degree of surface compliance critically influences sensory reweighting. Varying stiffness levels modulate the extent to which proprioceptive input is down-weighted (Schut et al., 2017), potentially altering the temporal dynamics of postural responses and contributing to the lower reliability observed for T_d_.

Several limitations should be considered when interpreting these findings. First, our experimental setup did not allow for the independent manipulation or isolation of proprioceptive and vestibular inputs. Therefore, we cannot determine whether the observed increase in visual weighting among older adults reflects a reduction in proprioceptive function, vestibular function, or both. Second, the study involved healthy older adults, which may limit the generalizability of the results to individuals with greater functional impairments or clinical balance deficits. One participant in the younger age group was excluded from the analysis due to behavior that deviated markedly form the rest of the sample. However, the data and model fit appeared good, reflecting very large sway responses to the visual scene movements. The participant’s disproportionately high reliance on visual input for spatial orientation may reflect an individual sensory integration deficit. To strengthen the robustness of our findings, future studies should include a broader range of participants across the adult lifespan rather than relying solely on two distinct age groups and also include more frail and less active participants. This would help clarify whether the observed patterns represent a gradual age-related trend.

In conclusion, this study provides evidence that visual dependence in balance control increases with age. The novelty is the use of sophisticated analysis methods that allowed for the identification of sensory contributions to balance control itself, rather than using indirect methods. While older adults showed significantly higher visual weighting, no age-related differences were observed in other core model parameters, including feedback gains and neural time delay, suggesting that the fundamental motor control mechanisms remain largely preserved in healthy aging. These findings underscore the potential of targeting sensory integration, particularly visual dependence, in balance training and fall prevention strategies for older adults.

